# Leading and Following: Noise Differently Affects Semantic and Acoustic Processing during Naturalistic Speech Comprehension

**DOI:** 10.1101/2023.02.26.529776

**Authors:** Xinmiao Zhang, Jiawei Li, Zhuoran Li, Bo Hong, Tongxiang Diao, Xin Ma, Guido Nolte, Andreas K. Engel, Dan Zhang

## Abstract

Despite the distortion of speech signals caused by unavoidable noise in daily life, our ability to comprehend speech in noisy environments is relatively stable. However, the neural mechanisms underlying reliable speech-in-noise comprehension remain to be elucidated. The present study investigated the neural tracking of acoustic and semantic speech information during noisy naturalistic speech comprehension. Participants listened to narrative audio recordings mixed with spectrally matched stationary noise at three signal-to-ratio (SNR) levels (no noise, 3 dB, -3 dB), and 60-channel electroencephalography (EEG) signals were recorded. A temporal response function (TRF) method was employed to derive event-related-like responses to the continuous speech stream at both the acoustic and the semantic levels. Whereas the amplitude envelope of the naturalistic speech was taken as the acoustic feature, word entropy and word surprisal were extracted via the natural language processing method as two semantic features. Theta-band frontocentral TRF responses to the acoustic feature were observed at around 400 ms following speech fluctuation onset over all three SNR levels, and the response latencies were more delayed with increasing noise. Delta-band frontal TRF responses to the semantic feature of word entropy were observed at around 200 to 600 ms leading to speech fluctuation onset over all three SNR levels. The response latencies became more leading with increasing noise and were correlated with comprehension performance and perceived speech intelligibility. While the following responses to speech acoustics were consistent with previous studies, our study revealed the robustness of leading responses to speech semantics, which suggests a possible predictive mechanism at the semantic level for maintaining reliable speech comprehension in noisy environments.

**Highlights:**

1. Leading responses were observed in the semantic-level neural tracking, with more leading latencies as noise increased.
2. Following responses were observed in the acoustic-level neural tracking, with more delayed latencies as noise increased.
3. Semantic-level neural tracking is correlated with comprehension performance and perceived intelligibility.
4. Distinct frequency bands were involved in speech semantic and acoustic processing.

## 1 Introduction

Noise is an inevitable part of daily life, from car horns on the streets to background music at parties, and it presents a significant challenge to verbal communication. Reliable speech comprehension in noisy environments is crucial in various situations such as education or emergency service. Despite the distortion of auditory information, individuals with normal hearing can comprehend speech with ease. Understanding the adaptive neural mechanisms that enable robust speech-in-noise comprehension is essential for clinical intervention for hearing/language-impaired groups and for developing hearing-aid techniques.

Neurophysiological studies have revealed important insights into how noise affects speech processing. Using the event-related techniques, cortical auditory evoked potentials (CAEP) elicited by auditory and speech stimuli have been found to show delayed latencies and reduced amplitudes under adverse conditions, including the early component P1-N1-P2 complex related to primary sound processing (Billings et al., 2009, 2011), and the later components such as the N2 component related to phonological analysis (Billings et al., 2009; Martin & Stapells, 2005; Tomé et al., 2015; Whiting et al., 1998) and the P3 component related to speech discrimination (Kaplan-Neeman et al., 2006; Koerner et al., 2017; Martin & Stapells, 2005; Whiting et al., 1998). In recent years, studies have focused more on the neural tracking of continuous speeches. i.e., the alignment between neural activities and the quasi-rhythmic fluctuations of continuous speech (see reviews, Brodbeck & Simon, 2020; Ding & Simon, 2014; Giraud & Poeppel, 2012; Lakatos et al., 2019; Obleser & Kayser, 2019). Specific temporal dynamics of neural tracking can be described via system identification methods such as the temporal response function (TRF; Crosse et al., 2016, 2021) by relating neural signals with speech features such as acoustic envelope. Neural tracking has been found to remain stable under mild and moderate noise, and it is regarded as an essential tool for segregating speech from the noisy background (Ding & Simon, 2013). Nevertheless, the TRF-based studies have also reported delayed latencies and/or reduced amplitudes of the neural tracking in noisy conditions (Gillis, Decruy, et al., 2022; Mirkovic et al., 2019; Muncke et al., 2022; Zou et al., 2019), similar to previous event-related studies. These results suggest an impaired acoustic processing efficiency in noisy environments (Gillis, Decruy, et al., 2022; Kaplan-Neeman et al., 2006). In addition to auditory processing, semantic processing also plays a vital role in speech-in-noise comprehension and has been paid substantial emphasis.

Semantic processing could be a crucial factor in robust speech comprehension against noisy environments. Numerous research has shown that coherent semantic context enabling anticipating upcoming stimuli contributes to an effective understanding of degraded speech (Miller et al., 1951; Obleser & Kotz, 2010, 2011; Sohoglu et al., 2012; Zekveld et al., 2011). For example, Miller et al. (1951) found that words in coherent sentences had higher intelligibility compared with the same words in unrelated word lists during speech-in-noise comprehension. Regarding the influence of noise on semantic processing, such as the N400 component (Kutas & Federmeier, 2011; Kutas & Hillyard, 1984), several studies have reported robust or increased amplitude of N400 under mild degradation, which might be related to additional cognitive effort (Jamison et al., 2016; Romei et al., 2011; Zendel et al., 2015), while other studies reported reduced/delayed N400 for degraded speech, which might be related to damaged signal quality (Aydelott et al., 2006; Connolly et al., 1992; Daltrozzo et al., 2012; Obleser & Kotz, 2011; Strauß et al., 2013). These mixed results provided valuable information on the complex relationship between noise and semantic processing. Moreover, it was discovered that semantic processing includes early responses before the onset of the stimulus, which was considered to be associated with semantic prediction (Grisoni et al., 2017, 2021; Pulvermüller & Grisoni, 2020). Nevertheless, it is still unknown how this pre-onset response is modulated by noise at various signal-to-ratios (SNRs). These inconsistent results and inadequate explorations of noise effect on semantic processing may be due to limitations inherent in the event-related design. This design typically uses highly-controlled and short-duration speech units, such as individual words (e.g., Romei et al., 2011) or disconnected sentences (e.g., Strauß et al., 2013), which only contain limited semantic/contextual information.

The recent rise of the naturalistic speech paradigm is expected to expand our knowledge of the neural mechanisms of semantic processing during speech-in-noise comprehension (Z. Li & Zhang, 2023). Compared to the highly-controlled and short-duration speech units, continuous naturalistic speech stimuli provide a better resemblance to our daily communications because of a longer duration, more flexible content, and less deliberate semantic violations (Alday, 2019; Alexandrou et al., 2020; Hartley & Poeppel, 2020; Sonkusare et al., 2019; Willems et al., 2020; Wöstmann et al., 2017). Most of all, the continuous naturalistic speech stimuli provide rich context-based semantic information (Alday, 2019; Alexandrou et al., 2020; Hamilton & Huth, 2020; Sonkusare et al., 2019), which is indispensable for semantic prediction and reliable speech comprehension in chaotic daily environments. In addition, via state-of-art computational linguistic models, the semantic information of naturalistic speech can be quantified, and the semantic-level neural tracking can be directly measured (Broderick et al., 2018, 2019, 2021; Gillis et al., 2021; Koskinen et al., 2020; Mesik et al., 2021; Weissbart et al., 2020), presenting a powerful tool to investigating how semantic processing is affected by noise at different SNRs.

The two frequently adopted semantic features in speech-related neuroscience research are entropy and surprisal derived from information theory (Brodbeck et al., 2022; Donhauser & Baillet, 2020; Goldstein et al., 2022; Heilbron et al., 2022), which respectively measures the semantic uncertainty of the upcoming stimuli and the unexpectedness of the current stimulus (Pickering & Gambi, 2018; Willems et al., 2016). The word surprisal was found to be associated with the superior temporal gyrus and inferior frontal sulcus, etc. (Willems et al., 2016), and is linked to an N400-like neural response, i.e., negativity at around 400 ms within the central-parietal electrodes (Broderick et al., 2021; Gillis et al., 2021; Heilbron et al., 2022). The word entropy was associated with neural activities within the left ventral premotor cortex, left middle frontal gyrus and right inferior frontal gyrus, etc. (Willems et al., 2016). Furthermore, Goldstein et al. (2022) derived word entropy from deep language models (GPT-2) and correlated them with electrocorticography (ECoG) signals. The results indicated that entropy was related to neural activities in the left-lateralized channels at several hundred milliseconds before the word onset. This pre-onset response is consistent with the semantic prediction potential (SPP) in event-related studies as a direct neural signature for semantic prediction (Grisoni et al., 2021; Pulvermüller & Grisoni, 2020). A recent study by Yasmin et al. (2023) discovered that the N400-like response in semantic-level neural tracking remained robust under mild and moderate noise conditions and declined abruptly at the high-noise level (SNR = -3 dB). However, the noise effect on the pre-onset response in semantic-level neural tracking is still unexplored.

The current study aimed to investigate the neural mechanisms of speech-in-noise comprehension by simultaneously focusing on both the acoustic and semantic levels as well as both the pre-onset and the post-onset stages. A naturalistic speech comprehension paradigm was employed, as the naturalistic speech stimuli were expected to provide better ecological validity and contextual information (Alday, 2019; Sonkusare et al., 2019). 60-channel EEGs were recorded while the participants listened to spoken narratives at three SNRs (no noise, 3 dB, -3 dB). Following previous studies, the amplitude envelopes of the speech stimuli were extracted as the acoustic feature (Di Liberto et al., 2015; O’Sullivan et al., 2015). Two typical semantic features were calculated by a Chinese NLP model, i.e., word entropy and word surprisal (Gillis et al., 2021; Weissbart et al., 2020; Willems et al., 2016; Koskinen et al., 2020; Mesik et al., 2021; Broderick et al., 2021). The neural responses to the acoustic and semantic features were estimated using the TRF method (Crosse et al., 2016), which yields the spatiotemporal dynamics of how our brain tracks these features in naturalistic speeches. The pre-onset and post-onset responses in the current study were defined as significant TRF responses with negative and positive time lags, respectively. Especially, we conducted TRF analyses and detected significant TRF responses separately at different SNR levels to capture all potential neural signatures. We hypothesize that the acoustic-level TRF could be related to delayed peak latencies or reduced amplitudes under noisy conditions as in previous studies (Gillis, Decruy, et al., 2022; Mirkovic et al., 2019; Muncke et al., 2022; Zou et al., 2019). As for the semantic-level TRF, we hypothesize that both the pre-onset and post-onset response could show resilience against noise (Yasmin et al., 2023). By exploring the pre-onset and post-onset temporal dynamics of low- and high-level processing, this study hopes to gain a more complete overview of the noise effect on neural processing during naturalistic speech comprehension.

## 2 Methods

### 2.1 Participants

Twenty college students (10 females, ages ranging from 19 to 28 years old) participated in the study. The sample size was determined to be sufficient by reference to previous TRF-based studies on the human speech processing (e.g., Broderick et al., 2018; Di Liberto et al., 2015). One male participant was excluded due to technical problems during data recording. The data of the remaining nineteen participants (age: mean ± SD = 21.79 ± 1.99) were included in the subsequent analyses. All participants were native Chinese speakers, right-handed, with normal hearing and normal or corrected-to-normal vision by self-report. The study was conducted in accordance with the Declaration of Helsinki and was approved by the local Ethics Committee of Tsinghua University. Written informed consent was obtained from all participants.

### 2.2 Materials

Thirty narrative audio recordings from our previous studies (Z. Li et al., 2021, 2022) were used as stimuli. These audio recordings were recorded from six native Chinese speakers with professional training in broadcasting. The participants were unfamiliar with the content of these narrative audio recordings, which were about speakers’ personal experiences on daily-life topics adapted from the National Mandarin Proficiency Test. Each narrative audio recording lasted for around 90 s and was recorded by a regular microphone at a sampling rate of 44,100 Hz in a sound-attenuated room.

These speech stimuli were further processed into three versions at three different SNR levels: no-noise (NN), low-noise (SNR = 3 dB), and high-noise (SNR = -3 dB), where speech intensity percentage was 100%, 60%, and 40%, respectively. This procedure was achieved by adding spectrally matched stationary noise, which was generated based on a 50th-order linear predictive coding (LPC) model estimated from the original speech recording (Broderick et al., 2018). The SNR levels were selected following previous studies (Ding & Simon, 2013), and were produced by varying the noise intensity while keeping the intensity of original speech (measured by its root mean square) constant (Ding & Simon, 2013).

For each narrative audio recording, two four-choice questions were prepared by the experimenters to assess one’s speech comprehension performance. These questions and the corresponding choices were targeted at detailed narrative contents that would demand significant attentional efforts. For instance, one question following a narrative audio recording about one’s major was, “What is the speaker’s most likely major as a graduate student? (说话人的研究生专业最可能是什么?)” and the four choices were 1) Social science, 2) International politics, 3) Pedagogy and 4) Psychology (1. 社会科学, 2. 国际政治, 3. 教育学 and 4. 心理学).

### 2.3 Procedure

Before the start of the experiment, the participants had one practice trial to get familiar with the procedure, with an additional narrative audio recording at the no-noise level not used in the formal experiment. The formal experiment consisted of 30 trials, with 10 trials per SNR level. In each trial, the participants listened to narrative audio recordings at one of the three SNR levels. The participants were required to maintain visual fixation on a fixation cross displayed on the computer screen in front of them and to minimize eye blinks and all other motor activities during listening. The order of the narrative audio recordings and their assigned SNR levels was randomized for each participant.

After each trial, the participants were instructed to answer two four-choice questions about the content of the narrative audio recording using the computer keyboard. The averaged accuracies across all trials (separately for each SNR level) were used to reflect the participants’ comprehension performance. After completing these questions, the participants were instructed to rate the perceived clarity and intelligibility of the narrative audio recording on a 100-point rating scale and rested for at least 5 s before moving on to the next trial. No feedback was given to the participants about their comprehension performance during the experiment.

The experimental procedure was programmed in MATLAB using the Psychophysics Toolbox 3.0 (Brainard, 1997). The speech stimuli were delivered to listeners seated in a sound-attenuated room via an air-tube earphone (Etymotic ER2, Etymotic Research, Elk Grove Village, IL, USA) to avoid environmental noise and equipment electromagnetic interference. The volume of the audio stimuli was adjusted individually for each participant to a comfortable level, and it was kept consistent across trials. The experimental procedure is illustrated in Figure 1.

**Figure 1.**
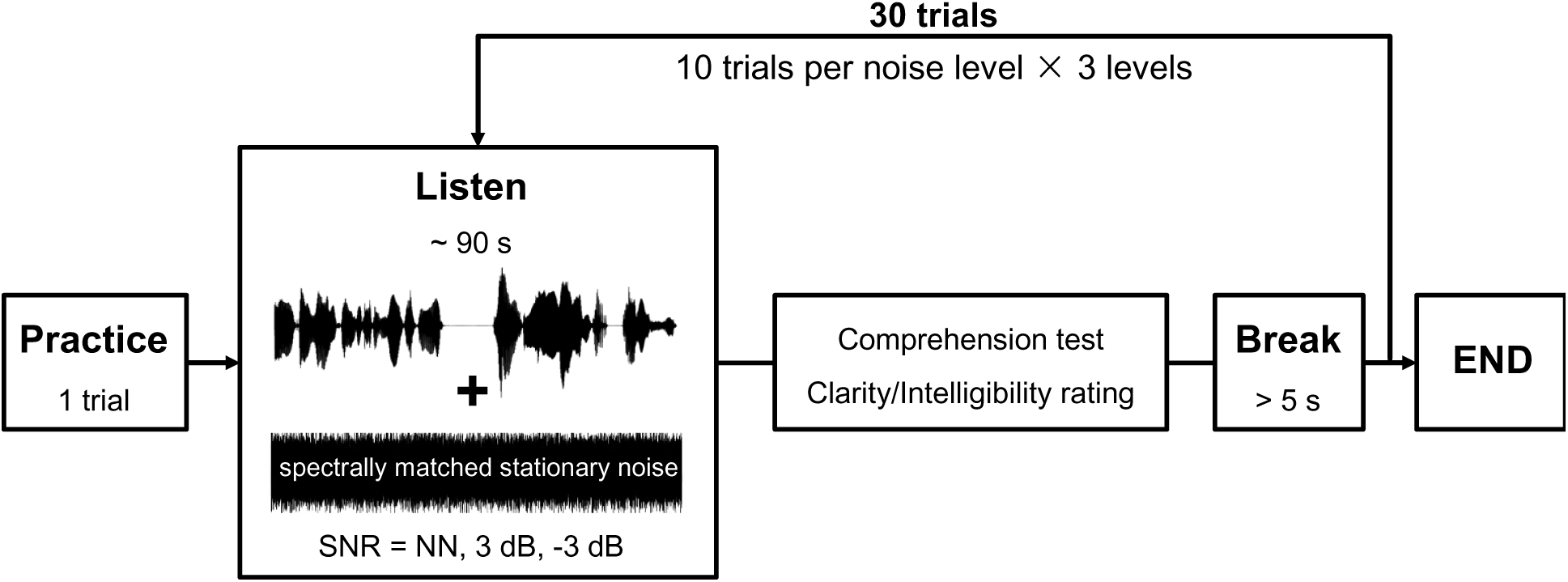
Experimental procedure. The participants listened to 30 naturalistic narrative audio recordings which each lasted around 90 s. These audio recordings were mixed with three levels of spectrally matched stationary noise: no-noise (NN), low-noise (SNR = 3 dB), and high-noise (SNR = -3 dB). The 60-channel EEG signals were recorded during listening. After listening to each narrative audio recording, the participants were required to complete a comprehension test and report the clarity and intelligibility rating. In the comprehension test, two four-choice questions per audio recording based on the narrative content were used.

### 2.4 EEG recording and preprocessing

EEG signals were recorded from 60 channels with a NeuroScan amplifier (SynAmp II, NeuroScan, Compumedics, USA) at a sampling rate of 1000 Hz. Electrodes were positioned according to the international 10–20 system, including FP1/2, FPZ, AF3/4, F7/8, F5/6, F3/4, F1/2, FZ, FT7/8, FC5/6, FC3/4, FC1/2, FCZ, T7/8, C5/6, C3/4, C1/2, CZ, TP7/8, CP5/6, CP3/4, CP1/2, CPZ, P7/8, P5/6, P3/4, P1/2, PZ, PO7/8, PO5/6, PO3/4, POZ, Oz, O1/2, referenced to an electrode between CZ and CPZ with a forehead ground at FZ. Electrode impedances were kept below 10 kOhm for all electrodes throughout the experiment.

The recorded EEG data were first notch filtered to remove the 50 Hz powerline noise. Independent Component Analysis (ICA) was performed to remove artifacts such as eye blinks and eye movements based on visual inspection. Around 4–12 independent components (ICs; mean = 6.6) were removed per participant. The remaining ICs were then back-projected onto the scalp EEG channels to reconstruct the artifact-free EEG signals. The EEG signals were then re-referenced to the average of all scalp channels and downsampled to 128 Hz. Afterward, EEG signals were filtered into the delta (1–4 Hz) and theta (4–8 Hz) bands, which have been previously reported to be important for speech neural tracking (Ding et al., 2014; Etard & Reichenbach, 2019; Keitel et al., 2017; Koskinen et al., 2020; J. Li et al., 2023; Peelle et al., 2013). For comprehensiveness, we also included the alpha (8−12 Hz) and beta (12−30 Hz) bands into analyses. Causal FIR (Finite Impulse Response) filters were employed to ensure that the filtered EEG signals were determined only by the current and previous data samples, which was important for the present study focusing on the fine-grained time course, particularly considering the pre-onset neural responses (de Cheveigné& Nelken, 2019).

These preprocessed EEG signals were segmented into 30 trials, from 5 to 90 s (duration = 85 s), relative to the speech onsets of each trial to avoid possible onset and offset effects. All preprocessing was conducted offline using MATLAB and the Fieldtrip toolbox (Oostenveld et al., 2011).

### 2.5 Feature extraction

Three types of features were extracted to represent the acoustic (amplitude envelope) and semantic (word entropy, word surprisal) information for each narrative audio recording. An example of these speech features is illustrated in Figure 2A.

**Figure 2.**
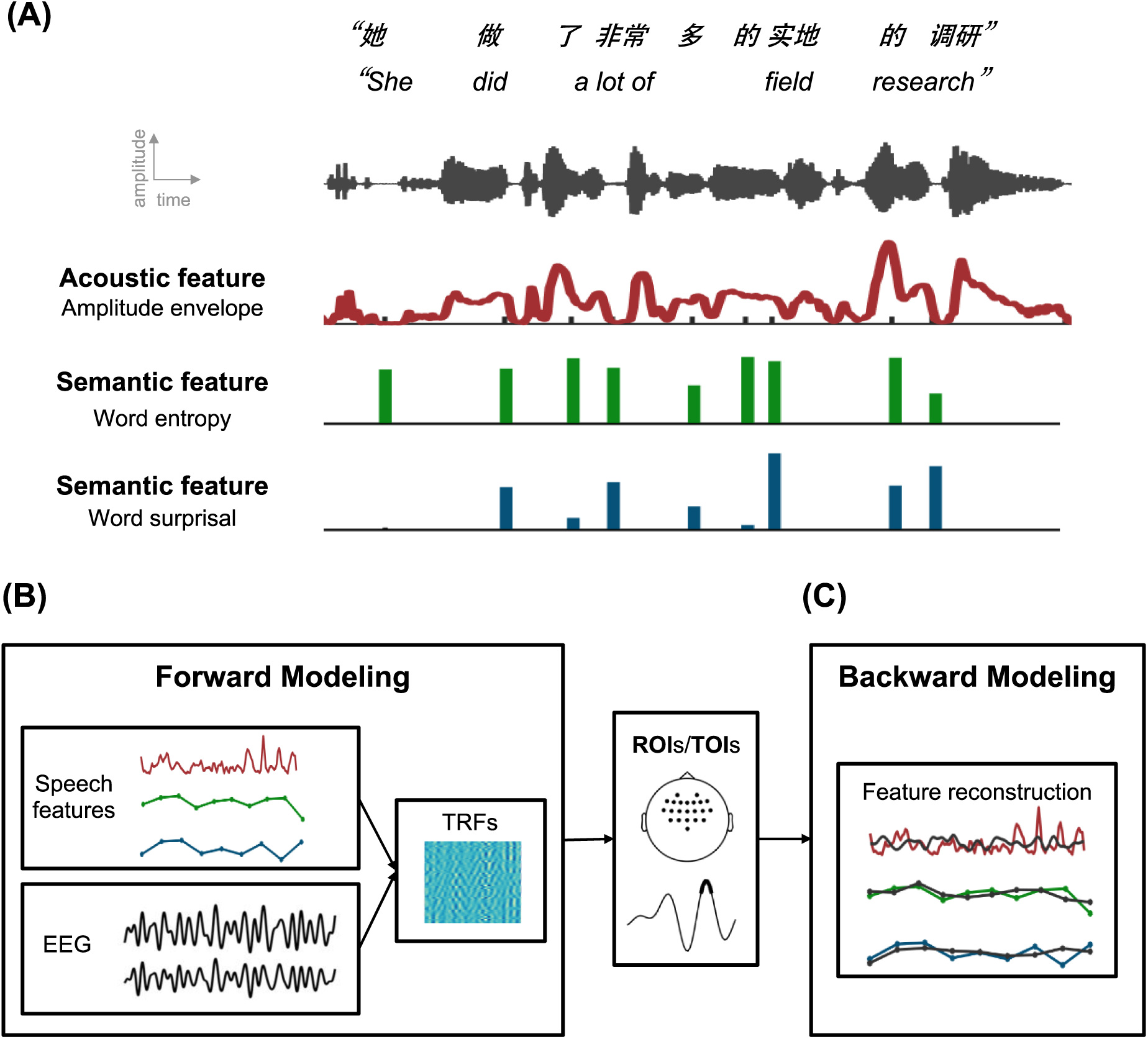
Speech feature extraction and Temporal Response Function analyses. (A) Three types of speech features were extracted, including one acoustic feature (amplitude envelope) and two semantic features (word entropy and word surprisal). The two semantic features were derived from a computational linguistic model and one-dimensional vectors were generated with impulses manipulated with semantic feature values at the corresponding word onsets time. (B) Forward modeling. TRFs were extracted by regressing each of the three types of speech features against the EEG signals separately. The significance of these TRFs was estimated by comparing them with the corresponding control TRFs, which were modeled based on EEG signals and shuffled speech features. The resulting spatiotemporal ranges were identified as regions of interest (ROIs) and time lags of interest (TOIs). (C) Backward modeling. These three types of speech features were separately reconstructed through backward TRF models, and the reconstruction accuracy (Pearson’s *r*) depicted the strength of neural tracking. Control backward models were constructed with EEG signals and shuffled features.

#### Acoustic feature

The amplitude envelope for each narrative audio recording was calculated as the absolute values after a Hilbert transform and then downsampled to the sampling rate of 128 Hz to match that of the EEG signals.

#### Semantic features

Before feature extraction, the narrative audio recordings were converted to text by Iflyrec software (Iflytek Co., Ltd, Hefei, Anhui) and then segmented into words based on the THU Lexical Analyzer for Chinese (THULAC) toolbox (Sun et al., 2016).

Two semantic features, word entropy and word surprisal, were extracted. Word entropy measures the uncertainty of predicting the upcoming word based on the context so far and was calculated as equation (1):

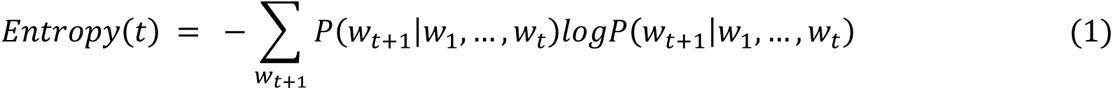

Word surprisal measures how surprising the current word is given the previously encountered words and was calculated as equation (2):

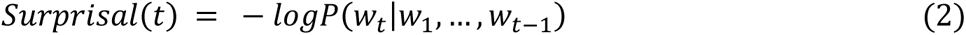

Where 𝑤_1_, …, 𝑤_𝑡−1_ are the existing word sequence and 𝑃(𝑤_𝑡_|𝑤_1_, …, 𝑤_𝑡−1_) is the conditional probability of next word (Willems et al., 2016). These NLP calculations were conducted by ADAM, a widely accepted Long-Short Term Memory (LSTM) Neural Network model (Kingma & Ba, 2015). The model was trained on the corpora corpus of the People’s Daily with 534,246 Chinese words. See Supplementary Table S1 for more information about the model and Supplementary Table S2 for more information about the descriptive statistics of semantic features.

After extracting the semantic features of each word, the word onset timings were estimated via Iflyrec software. Impulses at the word onset time were manipulated with corresponding semantic feature values to generate one-dimensional “semantic vectors” (e.g., Broderick et al., 2018; Gillis et al., 2021). The sampling rate of the semantic vectors was 128 Hz to match the EEG signals.

### 2.6 Modeling of the stimulus-response relationship

The Temporal Response Function (TRF) modeling method based on ridge regression was adopted to explore the relationship between the neural activities and the three types of stimulus features (Crosse et al., 2016, 2021). Forward modeling was first used to illustrate the specific spatiotemporal response patterns and identify key electrodes and time lags in TRF responses of the corresponding speech feature, and then backward modeling was adopted to verify the possible contribution of these identified neural correlates (e.g., Broderick et al., 2019; Etard & Reichenbach, 2019). The overall procedure of the modeling analyses is shown in Figure 2B and 2C.

#### Forward modeling

With a forward modeling approach, we described neural response patterns to different speech features by linear spatiotemporal filters called TRFs, which measure how neural signals from different regions are modulated by stimulus features at different time lags (Crosse et al., 2016). The estimated TRF together with the corresponding speech feature was used to predict the EEG responses from each electrode. The prediction accuracy measured as the Pearson’s correlation between the actual and predicted EEG signals represents the performance of the forward model. The TRF, *w*, is measured by equation (3):

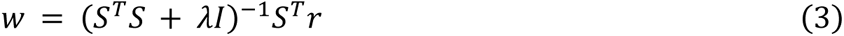

Where *S* is the lagged time series of the stimulus features, *r* is the neural signals, and *I* is the identity matrix. The time lags for forward modeling were chosen to cover a relatively broad time range, from -1000 to 1000 ms (Goldstein et al., 2022; J. Li et al., 2021, 2023). The *λ* is the regularization parameter used to prevent overfitting and ranged between 0.1 and 1000 in steps of the powers of 10 empirically (Gillis, Van Canneyt, et al., 2022). The cross-validation procedure was implemented in a leave-one-trial-out procedure within each participant: each time, the model was trained based on 9 trials and tested on the left-out trial, which was repeated for each of the 10 trials at three SNR levels separately. The *λ* value that produced the highest prediction accuracy averaged across trials after cross-validation was selected as the regularization parameter for all trials at a certain SNR level per participant. TRF amplitudes were further transformed into *z*-scores before statistical analyses (Ding et al., 2014; Gillis et al., 2021; J. Li et al., 2021).

The statistical significance of the estimated TRFs was estimated by constructing control TRF models (Weissbart et al., 2020). We built control models by constructing TRF models using shuffled stimulus features and the EEG recordings in the same way as for the computation of the actual TRFs. The shuffled amplitude envelope was constructed by randomly shuffling the feature value within a trial while keeping the timing of the quiet fragments. The shuffled word entropy and word surprisal were constructed by randomly shuffling the feature value within a trial while keeping the timing of impulses. Therefore the speech features that described acoustic and linguistic word onsets were not altered and thus left no impact on TRFs’ significance (Weissbart et al., 2020). The shuffling was repeated 1,000 times and resulted in 1,000 control TRFs for a corresponding actual TRF.

A nonparametric cluster-based permutation test was applied to account for multiple comparisons (Maris & Oostenveld, 2007). For each electrode-time bin in the actual and control TRFs, a one-sample *t*-test was used to examine whether the TRF amplitudes significantly differed from 0. Then neighboring electrode-time bins with an uncorrected *p*-value less than 0.01 were combined into clusters. The minimum number of neighboring significant channels that was required for inclusion in a cluster was 2. For each cluster, the sum of the *t*-statistics was obtained. A null distribution was created from the 1,000 control test statistics, i.e., the maximum cluster-level *t*-statistics. The corrected *p*-value for each cluster was calculated as the proportion of control test statistics greater than the actual cluster-level *t*-statistics. Clusters with *p*-values below 0.05 were selected for further analyses. We implemented the same statistical analyses procedure for each of the 18 TRFs (3 stimulus features × 2 frequency bands × 3 SNR levels). The EEG electrodes and time lags from significant clusters were regarded as ROIs/TOIs. Then peaks were estimated within these ROIs/TOIs, and the peak amplitudes and peak latencies were compared across different SNRs.

#### Backward modeling

With a backward modeling approach, we simultaneously took neural signals from several electrodes to reconstruct stimulus features with a decoder. The reconstruction accuracy was measured as the Pearson’s correlation between the actual and reconstructed stimulus features. The decoder, *g,* is calculated by equation (4), where *R* is the lagged time series of the EEG data. The reconstructed feature, 𝑠^(𝑡), is calculated by equation (5) where *n* is the EEG electrodes, and 𝜏 is the time lags (Broderick et al., 2019).

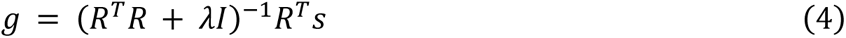

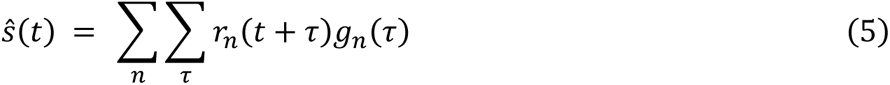

Only the exact ROIs/TOIs from significant clusters found in the forward modeling were included in the backward modeling. The EEG signals and stimulus features were downsampled to 64 Hz for better computational efficiency. The same leave-one-trial-out cross-validation procedure as in the forward modeling approach was conducted to obtain the optimal regularization parameter and calculate the reconstruction accuracy. We also estimated the control decoders using the same shuffling methods in forward modeling. The reconstruction accuracies from the 1,000 control decoders were averaged and compared with the actual decoder via a one-tailed paired *t*-test, and the *p* values of clusters were corrected via the false discovery rate (FDR) method (Benjamini & Hochberg, 1995).

In sum, the analyses of neural tracking followed two main steps. (1) We calculated the actual forward model and control forward models and identified ROIs/TOIs according to clusters with significant differences between them. (2) We estimated the reconstruction accuracy based on these ROIs/TOIs. This procedure resulted in (1) the specific spatiotemporal TRF response pattern and (2) the strength of neural tracking (reconstruction accuracy) for analyses.

Given that the neural signatures of speech processing could exhibit different spatiotemporal patterns at various SNR levels (e.g., Bidelman & Howell, 2016; Billings et al., 2009; Strauß et al., 2013), we conducted separate statistical tests for identifying significant clusters in the TRF responses at different SNR levels, in order to capture all potentially significant results without missing anything.

We classified these significant clusters into two types based on their spatiotemporal dynamics: those with largely overlapped spatiotemporal patterns across all SNR levels, which could represent a reliable response across all SNR levels, and those with unique patterns at a certain SNR level, which might signify distinct processing mechanisms under certain circumstances. This was achieved by visual inspection and calculating a similarity index, which derived from the average of the temporal and spatial similarity. See Supplementary Figure S2 for more information. The consistent clusters were compared to explore how the commonly shared neural signature adapted to noise, while the unique clusters received less attention in subsequent analyses. Linear-mixed effect (LME) models and Spearman’s correlation was conducted to examine the relationship between neural tracking and behavioral performance.

Before modeling, the three types of stimulus features across all trials and EEG signals across all channels were z-scored as recommended to ensure consistent scaling (Crosse et al., 2016, 2021). Modeling and analyses for different stimulus features were conducted independently. Considering we focused on the neural tracking of underlying hierarchical information in speech rather than physical stimulus, we adopted the same stimulus features of no-noise speech for the other two SNR levels (Ding & Simon, 2013; Fuglsang et al., 2017). The forward and backward modeling was conducted in MATLAB using the Multivariate Temporal Response Function (mTRF) toolbox (Crosse et al., 2016). The cluster-based permutation test was conducted in the FieldTrip toolbox (Oostenveld et al., 2011). Other statistical analyses were conducted via MATLAB functions and IBM SPSS Statistics software (IBM corp., 2019).

## 3 Results

### 3.1 Behavioral performance

The speech comprehension performance was measured as the averaged response accuracy of the four-choice questions and was found to be significantly different among the three SNR levels (rmANOVA, *F*(2, 36) = 6.74, *p* = .003). The speech comprehension performance was 95.26 ± 1.05%, 90.79 ± 2.10%, and 85.26 ±2.63% (mean ±SE) at the no-noise level, low-noise level, and high-noise level, respectively. The comprehension performance at the high-noise level was significantly lower than that at the no-noise level (post-hoc *t*-test, *p* = .006, Bonferroni corrected). In addition, it should be noted that the comprehension performance was still well above chance level even at the high-noise level (one-tailed *t*-test, *t*(18) = 22.88, *p* < .001).

The subjective ratings of clarity and intelligibility showed a similar pattern with significant differences among the SNR levels (rmANOVA, *F*(2, 36) = 148.32 and 35.31, *p*s < .001). The normalized clarity rating scores were 0.94 ±0.01, 0.66 ±0.04, and 0.35 ±0.04 (mean ±SE), and the normalized intelligibility rating scores were 0.93 ± 0.01, 0.88 ± 0.02, and 0.73 ± 0.03 (mean ± SE) at the no-noise, low-noise, and high-noise level, respectively. Post hoc *t*-tests revealed significant pairwise differences for all possible comparisons (*p*s < .01, Bonferroni corrected). The behavioral performance is illustrated in Figure 3. These results suggested that the effect of noise on speech comprehension and perception was effectively manipulated.

**Figure 3.**
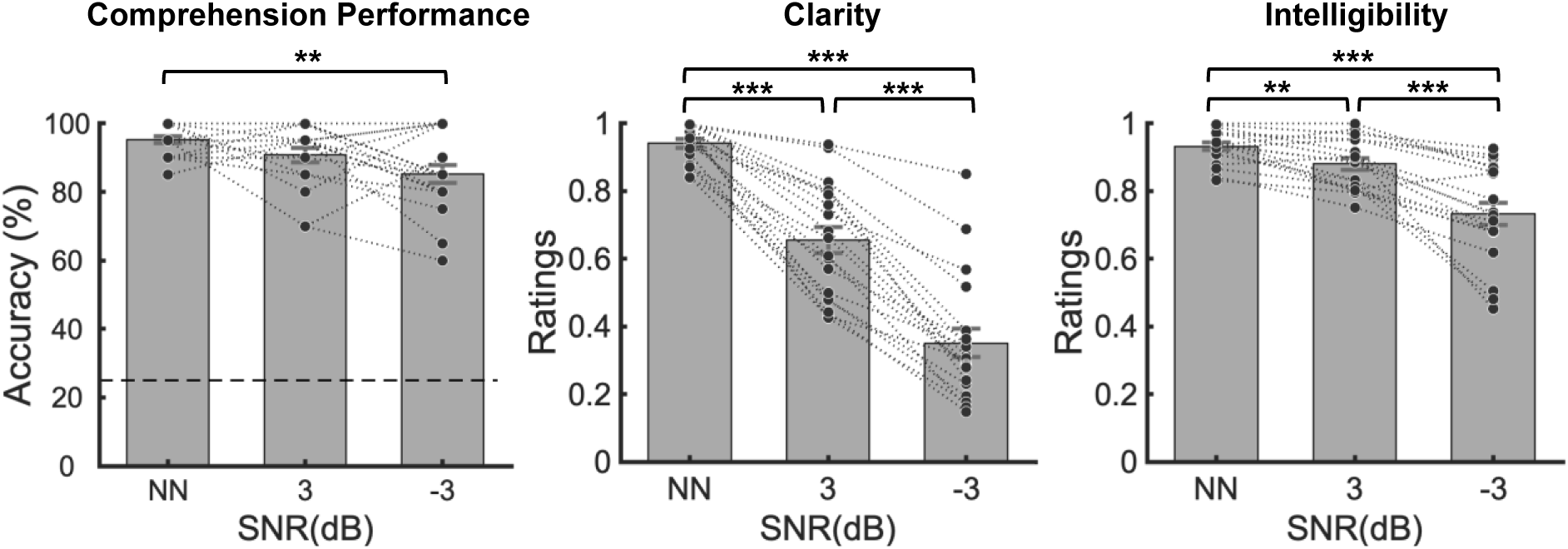
Behavioral results. Black dots indicate data points from each participant. Error bars denote the standard error. *: *p* < .05, **: *p* < .01, ***: *p* < .001.

### 3.2 Summary of all significant clusters in the acoustic- and semantic-level TRF responses

We summarized all significant clusters in the acoustic- and semantic-level TRF responses in Figure 4. Significant clusters were only found in the delta/theta bands but not alpha/beta bands (see Supplementary Figure S1 for more information). The specific time lags of TOIs of clusters were listed in Supplementary Table S3-4. According to visual inspection and the similarity index (shown in Supplementary Figure S2), these significant clusters were classified into responses that exhibited relative consistency across different SNR levels, as well as distinctive response at a certain SNR level.

**Figure 4.**
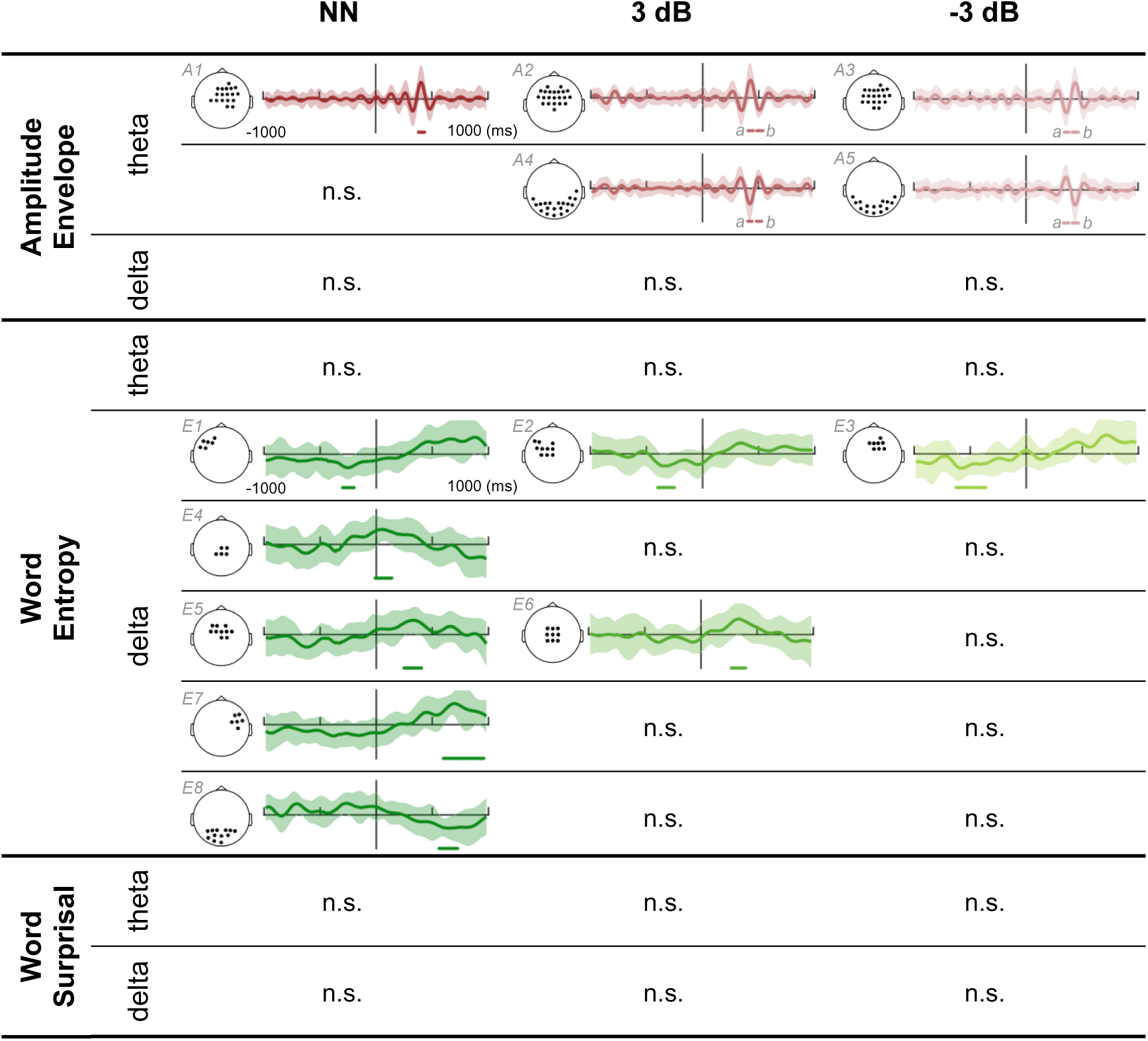
Significant clusters in TRF responses to (A) amplitude envelope, (B) word entropy, and (C) word surprisal at different SNR levels and frequency bands. Significant clusters are numbered A1−A5 and E1−E8. Clusters with similar spatiotemporal patterns are organized in the same row. ROIs of clusters are shown as black dots in the corresponding topography. The colored curves are the mean of TRFs averaged among ROIs across participants. The shaded areas denote the standard error of TRFs. The colored horizontal line below the TRF curve indicates the TOIs of the cluster. The *a* and *b* refer to clusters with the same ROIs but different TOIs. n.s.: no significant cluster was found.

Clusters with largely overlapped spatiotemporal patterns across all SNR levels were found in theta-band acoustic-level TRFs (i.e., A1, A2, A3) and delta-band entropy-based semantic-level TRFs (i.e., E1, E2, E3). Detailed analyses of them are demonstrated in the section 3.3 and 3.4, respectively. Several clusters with similar spatiotemporal patterns shared by certain SNR levels, such as the acoustic-level TRF response within the occipital electrodes (i.e., A4 and A5) and the post-onset entropy-based semantic-level TRF responses within the central electrodes (i.e., E5 and E6). Analyses of them are demonstrated in Supplementary Figure S3 and S4. No further analysis was done for the other unique clusters at the no-noise level. No significant TRF responses to word surprisal were found.

### 3.3 Acoustic-level TRF responses with delayed latencies as noise increases

Significant acoustic-level TRF responses in the theta band were found at all SNR levels and showed a similar positivity within central electrodes at around 300∼500 ms (i.e., A1, A2a, A3b), as demonstrated in Figure 5A. At the no-noise level, the TRF showed positivity in the central electrodes with a latency of around 400 ms (cluster-level *p* < .01). At the low-noise and high-noise levels, the TRF showed similar positivity in the central electrodes with a latency of around 430 (cluster-level *p* < .01) and 440 ms (cluster-level *p* < .01). We estimated the peak amplitude and peak latency for the positive peak at each SNR level. The peak latencies were significantly different among the three SNR levels (rmANOVA, *F*(2, 36) = 21.42, *p* < .001), and post-hoc *t*-tests revealed significantly delayed peak latencies as noise increased (*ps* < .05, Bonferroni corrected), as shown in Figure 5C. No significant differences were found in the peak amplitudes (rmANOVA, *F*(2, 36) = 0.64, *p* = .535).

**Figure 5.**
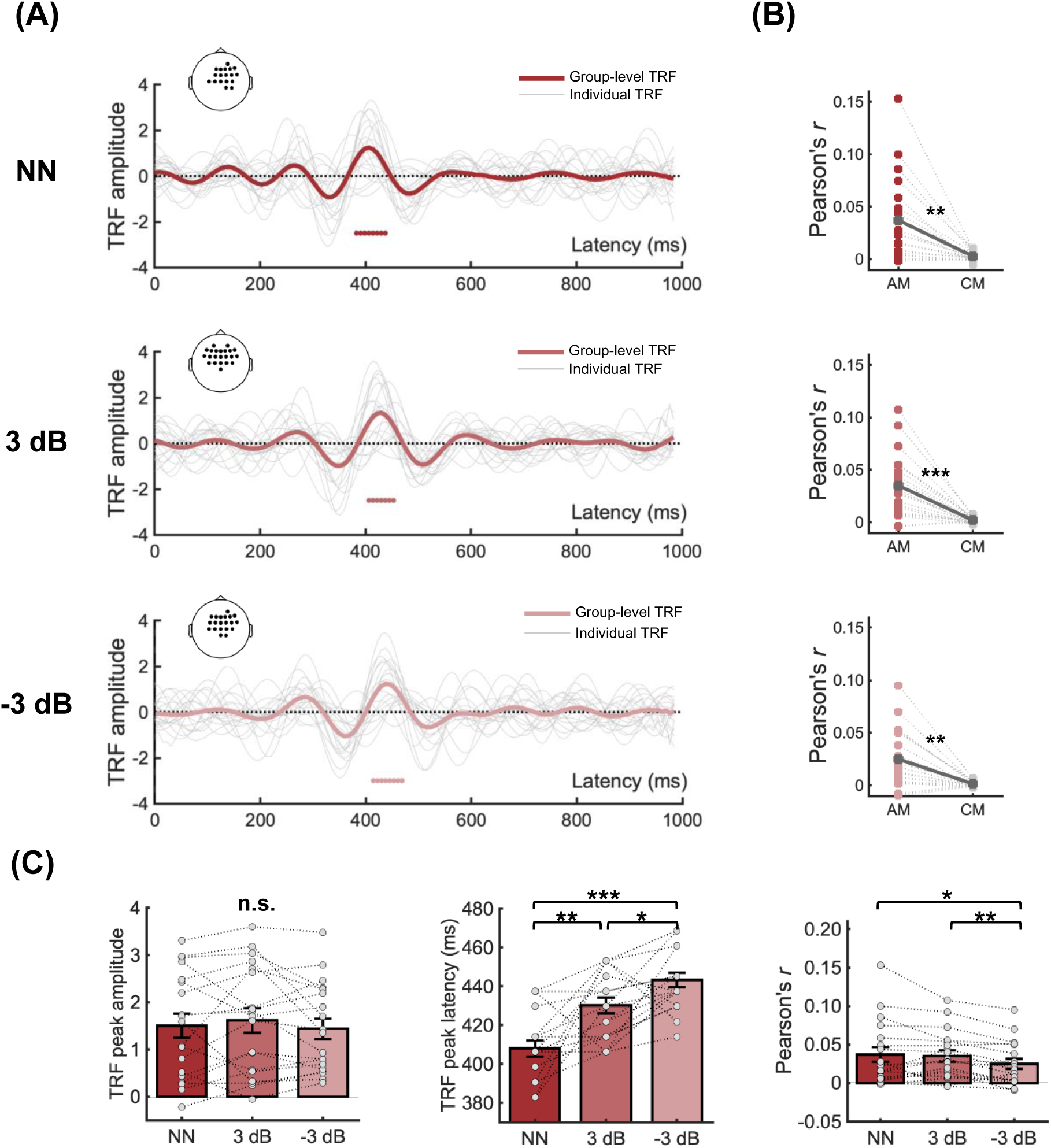
Acoustic-level TRF responses in the theta band at different SNR levels. (A) The bold curves in different shades of red are the mean of TRFs averaged among ROIs across participants at the three SNR levels. The grey curves are TRFs averaged among the ROIs of each participant. The colored horizontal line at the bottom of each plot indicates TOIs over which TRFs differed significantly from the control models. Dots in the corresponding topographies depicted the ROIs. (B) Reconstruction accuracy calculated from the ROIs/TOIs in (A). AM means actual models. CM means control models. (C) Noise effect on the peak amplitude, peak latency, and reconstruction accuracy. Grey dots indicate data points from each participant. Error bars denote the standard error. n.s.: not significant, *: *p* < .05, **: *p* < .01, ***: *p* < .001.

The reconstruction accuracies from the corresponding ROIs/TOIs were significantly higher in the actual decoders than in the control decoders (*p*s < .01, FDR corrected, Figure 5B). Comparing the reconstruction accuracies revealed significant differences among the three SNR levels (rmANOVA, *F*(2, 36) = 6.19, *p* = .010 with Greenhouse-Geisser correction), and post-hoc *t*-tests revealed significantly weakened neural tracking at the high-noise level compared with the no-noise level and the low-noise level (*p*s < .05, Bonferroni corrected, Figure 5C).

### 3.4 Semantic-level TRF responses with earlier latencies as noise increases

Significant semantic-level TRF responses to word entropy in the delta band were found at all SNR levels. They showed similar negativity at around 200∼600 ms leading to speech fluctuation onset, as demonstrated in Figure 6A. The time lags of pre-onset processing to word entropy showed a gradual advanced trend as noise increased. The time lag was approximately from around -300 ms to -180 ms at the no-noise level (cluster-level *p* < .05) and was from around -400 ms to -250 ms at the low-noise level (cluster-level *p* < .05), from around -630 ms to -360 ms at the high-noise level (cluster-level *p* < .01). We estimated the peak amplitude and peak latency for the pre-onset negative peak at each SNR level. The peak latencies were significantly different among the three SNR levels (rmANOVA, *F*(2, 36) = 58.08, *p* < .001), and post-hoc *t*-tests revealed that as noise increased the peak latencies were gradually earlier (*p*s < .05, Bonferroni corrected), as shown in Figure 6C. No significant differences were found in the peak amplitudes (rmANOVA, *F*(2, 36) = 0.63, *p* = .538).

**Figure 6.**
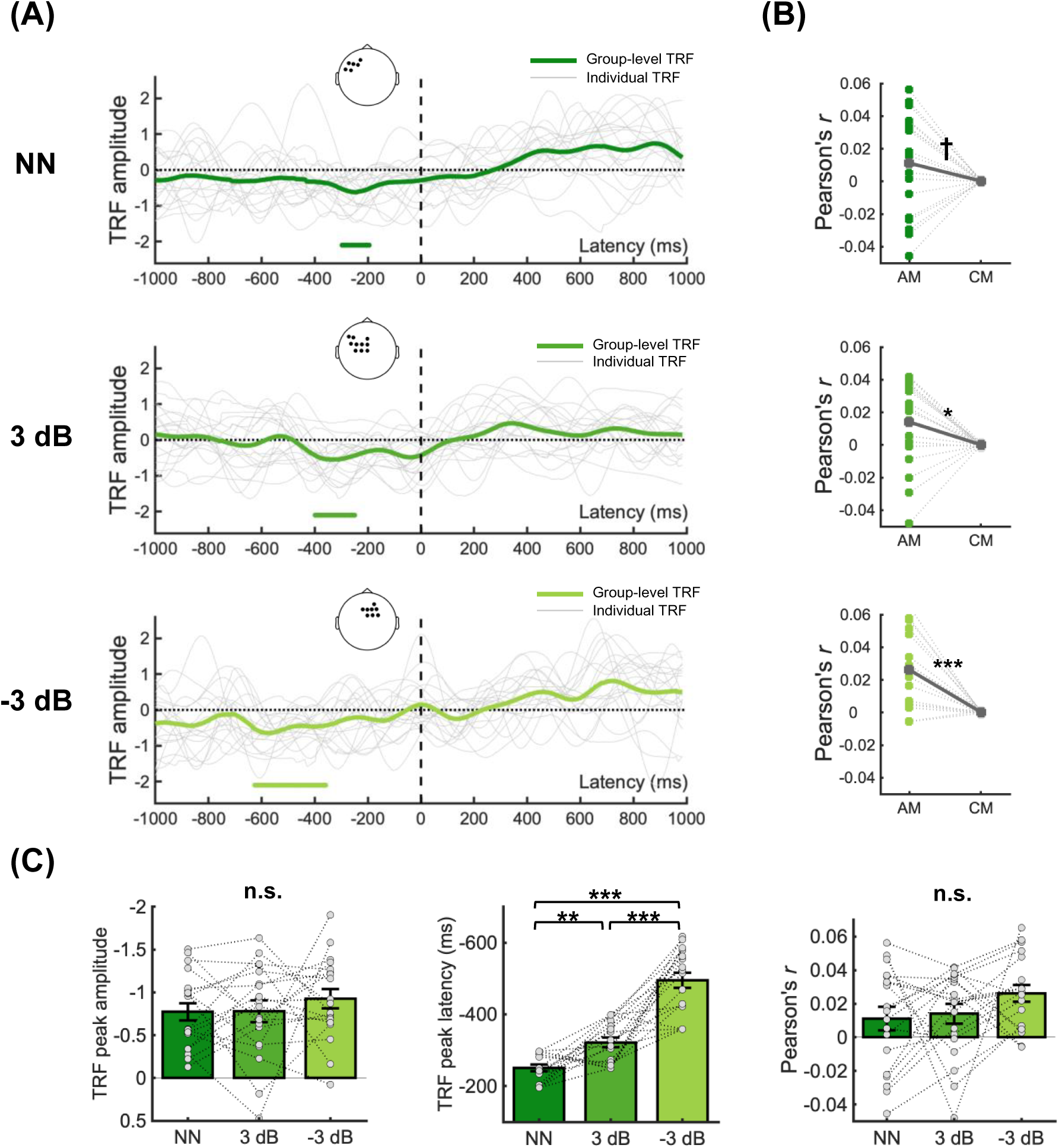
Semantic-level TRF responses to word entropy in the delta band at different SNR levels. (A) The bold curves in different shades of green are the mean of TRFs averaged among ROIs across participants at the three SNR levels. The grey curves are TRFs averaged among the ROIs of each participant. The colored horizontal line at the bottom of each plot indicates TOIs over which TRFs differed significantly from the control models. Dots in the corresponding topographies depicted the ROIs. (B) Reconstruction accuracy calculated from the ROIs/TOIs in (A). AM means actual models. CM means control models. (C) Noise effect on the peak amplitude, peak latency, and reconstruction accuracy. Grey dots indicate data points from each participant. Error bars denote the standard error. n.s.: not significant, †: *p* < .1, *: *p* < .05, **: *p* < .01, ***: *p* < .001.

The pre-onset TRF responses to word entropy exhibited different spatiotemporal patterns at three SNR levels. At the no-noise level, the ROIs included frontal-parietal electrodes and exhibited obvious left lateralization. At the low-noise level, the ROIs showed similar left-lateralized topological distribution but included more electrodes, while at the high-noise level, no obvious lateralization was observed in the frontal-parietal ROIs.

The reconstruction accuracies from corresponding ROIs/TOIs were significantly higher in the actual decoders than in the control decoders at both the low-noise level (*p* < .05, FDR corrected) and the high-noise level (*p* < .001, FDR corrected), but only marginally significant at the no-noise level (*p* = .073, FDR corrected), as shown in Figure 6B. Comparing the reconstruction accuracies among different SNR levels revealed no significant differences (rmANOVA, *F*(2, 36) = 1.64, *p* = .208, Figure 6C).

### 3.5 Correlation between TRF responses and behavioral performance

As the peak latencies of both the post-onset positive peak of acoustic-level TRFs and the pre-onset negative peak of semantic-level TRFs showed significant differences among SNR levels, we then created linear mixed effect models to explore whether these peak latencies were sensitive predictors of the behavioral performance with the following general formula:

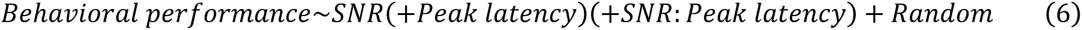

where “Behavioral performance” refers to the comprehension performance, clarity, or intelligibility ratings. “SNR” takes the values of speech intensity percentage, i.e., 100%, 60%, and 40%. “Peak latency” refers to the peak latencies of either the acoustic-level post-onset TRFs or the semantic-level pre-onset TRFs, depending on the specific model being investigated. “SNR:Peak latency” refers to the interaction between them. A random intercept per participant was included in the model. “Peak latency” and “SNR:Peak latency” were added between brackets to the general formula because these factors were included only if they led to a lower Akaike Information Criterion (AIC) which indicated a better fitting (Verschueren et al., 2022). Results showed that overall, the behavioral performance was correlated with SNR levels, which echoed the behavioral results in the section 3.1. More importantly, the earlier peak latencies of semantic-level pre-onset TRF response were correlated with the decreasing comprehension performance (LME, *β* = 2.61×10^-4^, *t*(52.64) = 1.853, *p* = .069) and the decreasing perceived intelligibility (LME, *β* = 9.26×10^-4^, *t*(40.48) = 3.497, *p* = .001). And the correlation with intelligibility was more prominent as noise increased (LME interaction, *β* = -7.69×10^-4^, *t*(45.80) = -1.987, *p* = .053), as illustrated in Table 1 and Figure 7. The relationships between the reconstruction accuracies and the behavioral performance were also examined through Spearman’s correlation and summarized in Supplementary Table S5 and Table S6.

**Figure 7.**
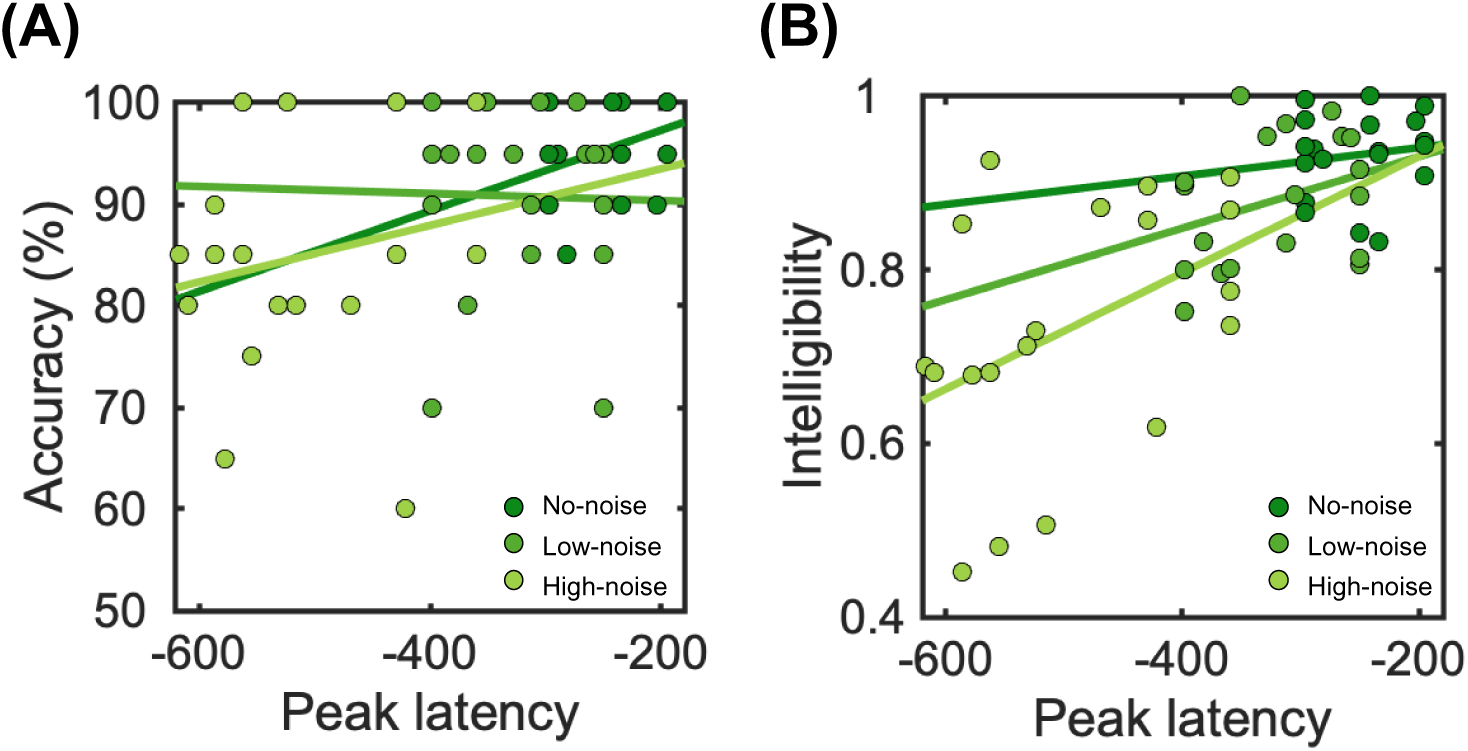
Correlation between the peak latencies of the semantic-level pre-onset TRFs and (A) the comprehension performance and (B) the perceived intelligibility. Colored dots indicate data points from each participant at different SNR levels.

**Table 1.** LME models of the behavioral performance as a function of SNR and peak latency. Each row indicates a different model. The SNR was given as a percentage (100%, 60%, 40%).

|  |  | SNR |  |  | Peak latency |  |  | SNR:Peak latency |  |  |
| --- | --- | --- | --- | --- | --- | --- | --- | --- | --- | --- |
| | | $\beta$ | $t$ | $p$ | $\beta$ | $t$ | $p$ | $\beta$ | $t$ | $p$ |
| Comprehension performance | TRF<sub>acoustic</sub> | $1.46 \times 10^{-2}$ | 3.835 | .001 | No lower AIC | | | No lower AIC | | |
| | TRF<sub>semantic</sub> | $6.23 \times 10^{-2}$ | 1.056 | .297 | $2.61 \times 10^{-4}$ | 1.853 | .069 | No lower AIC | | |
| Clarity | TRF<sub>acoustic</sub> | $8.87 \times 10^{-1}$ | 16.481 | < .001 | No lower AIC | | | No lower AIC | | |
| | TRF<sub>semantic</sub> | $8.87 \times 10^{-1}$ | 16.481 | < .001 | No lower AIC | | | No lower AIC | | |
| Intelligibility | TRF<sub>acoustic</sub> | $1.74 \times 10^{-1}$ | 6.050 | < .001 | No lower AIC | | | No lower AIC | | |
| | TRF<sub>semantic</sub> | $-1.34 \times 10^{-1}$ | -1.147 | .257 | $9.26 \times 10^{-4}$ | 3.497 | .001 | $-7.69 \times 10^{-4}$ | -1.987 | .053 |

## 4 Discussion

The current study investigated the neural tracking of hierarchical features of naturalistic speech in noisy situations using a TRF-based technique. Significant post-onset acoustic-level TRF responses were found within the central electrodes at around 400 ms, and the peak latencies were delayed as noise increased. Significant pre-onset semantic-level TRF responses were found within the frontal electrodes at around - 600∼-200 ms. The peak latencies showed a gradually advanced trend as noise increased, and increased advancement was correlated with decreasing comprehension performance and intelligibility. These findings indicated that noise differently modulates acoustic and semantic processing and suggested that robust and adaptive semantic pre-activation could play a vital role in reliable speech comprehension in noisy environments.

The delayed peak latency in the acoustic-level TRF responses as noise increased was in line with several previous studies, suggesting an impaired efficiency in challenging conditions with background noise (Gillis, Decruy, et al., 2022; Mirkovic et al., 2019; Muncke et al., 2022; Yasmin et al., 2023; Zou et al., 2019). As the frontally and centrally distributed channels (corresponding to the primary auditory cortex, superior temporal gyrus, premotor cortex, etc.) have been frequently reported to be related to the processing of speech acoustics (e.g., Bidelman & Howell, 2016; Broderick et al., 2019; Hickok & Poeppel, 2007; Zou et al., 2019), the present TRF results would imply similar recruitment of these brain regions for acoustic-level processing for naturalistic speech under various SNR levels. However, the post-onset 400-ms latency was later compared to the commonly reported latency of < 300 ms in previous studies (Gillis, Decruy, et al., 2022; Yasmin et al., 2023). This discrepancy could be due to the causal filter used for EEG signal preprocessing in the present study, possibly resulting in a delayed TRF compared to previous studies using noncausal filters, similar as reported by Etard and Reichenbach (2019). Alternatively, it could be possible that the latency modulation started earlier but only reached significance later for the present dataset, as the observed TRF responses exhibited an oscillatory pattern starting much earlier than 400 ms (Figure 5C). While an impaired processing efficiency has been associated with both amplitude and latency modulation by noise in previous studies (Muncke et al., 2022; Zion Golumbic et al., 2013; Zou et al., 2019), the present study together with a series of other studies reporting latency-only results would suggest latency as a more sensitive candidate for noisy speech processing (Ding & Simon, 2013; Kaplan-Neeman et al., 2006; Whiting et al., 1998).

At the semantic level, our findings on the pre-onset response to word entropy were consistent with recent studies, in which the neural responses to entropy have been reported to involve neural activities within the left hemisphere at up to 800 ms before onset (Goldstein et al., 2022; Weissbart et al., 2020; Willems et al., 2016). This pre-onset prediction mechanism for the upcoming stimuli was regarded as a fundamental computational principle in the human language processing (Goldstein et al., 2022). Our results echo these findings and highlight the potential of entropy as a promising index for exploring forward-looking prediction mechanisms.

More importantly, our results extend the present understanding of the predictive mechanism with the manipulation of the SNR levels and suggest a distinct mechanism for speech-in-noise comprehension at the semantic level. The significant pre-onset response to word entropy appeared at all SNR levels, which would imply that such a forward-looking prediction was robust against noise. In addition, we found that the peak latencies of the pre-onset responses became earlier with increasing noise, and that increased forward shift trend at each SNR level was correlated with poorer perceived intelligibility as well as decreasing comprehension performance. One possible explanation for this phenomenon is that our brain could adjust the timing of predictive processing in response to adverse environments. As semantic prediction can facilitate speech comprehension (Mattys et al., 2012; Miller et al., 1951; Obleser & Kotz, 2010; Pickering & Gambi, 2018; Zekveld et al., 2011), the brain relies on it more heavily as noise increases to counteract distorted auditory input. Nevertheless, noise can increase the processing load and decrease the processing efficiency (Gillis, Decruy, et al., 2022; Kaplan-Neeman et al., 2006; Kong et al., 2014; Mirkovic et al., 2019). To compensate for the interference, the neural system initiates the pre-onset response earlier and extends it for a longer duration, giving our brain more time for the preparation of the upcoming speech information. The more degraded the speech, the greater the need for this kind of “early-bird” compensation. Another possible explanation is that in noisy environments our brain relies more on longer-range prediction based on higher-level context information to enhance speech comprehension. According to a recent study based on GPT-2 and functional Magnetic Resonance Imaging (fMRI) (Caucheteux et al., 2023), the forward-looking prediction involved hierarchical representations and multiple time scales, with the maximum of forecast distance reaching 8 words (corresponding to approximately 3.15s). Future studies could employ local and context-unified entropy (e.g., Brodbeck et al., 2022) and longer time windows to further elucidate the noise effect on the forward-looking prediction. Overall, our findings suggest that the brain has a robust and adaptive prediction mechanism for reliable speech comprehension in noisy environments. As such a pre-onset signature was not observed at the acoustic level, our results suggest the predictive mechanism might be mainly focused at the semantic level (Goldstein et al., 2022; Grisoni et al., 2021), where the speech information is expected to be more abstract and more robust against noise (Yasmin et al., 2023).

Interestingly, the spatial patterns of TRF responses to word entropy showed left-lateralization at the no-and low-noise levels and recruited bilateral hemispheres at the high-noise level. The left-lateralization has been reported in studies on speech-in-noise comprehension (Z. Li et al., 2021) and was found to be sensitive to linguistic content (Peelle et al., 2013), word entropy (Willems et al., 2016), and semantic expectancy (Golestani et al., 2013; Obleser & Kotz, 2010). Our results would support the left-lateralized brain regions for predictive speech processing at the semantic level. Meanwhile, research has reported that regions within the right hemisphere, such as the right inferior frontal gyrus, are sensitive to semantic features such as entropy (Willems et al., 2016) and that the involvement of the right hemisphere increased under degraded conditions (Bidelman & Howell, 2016), which was hypothesized as the possible recruitment of additional regions for compensation (Shtyrov et al., 1998, 1999). Accordingly, our results suggest that the involvement of bilateral hemispheric in adverse environments might reflect a semantic-related compensation mechanism.

Our results suggested the specificity of the frequency band for processing different levels of speech information. Specifically, acoustic-level TRF response was primarily associated with the theta band whereas semantic-level TRF neural response was dominated by the delta band (Figure 4). This could be explained as that theta- and delta-band neural tracking have different functional roles: the former is related to acoustic processing while the latter is related to sematic/syntactic processing (Dai et al., 2022; Ding et al., 2014; Etard & Reichenbach, 2019; Kösem & van Wassenhove, 2017; J. Li et al., 2023). Alternatively, this distinction could be related to the intrinsic temporal properties of the speech features (Lalor, 2018), that is, a faster acoustic-level fluctuation at the theta rhythm and a slower semantic-level fluctuation at the word rate similar to the delta rhythm. Further studies could employ careful experimental manipulation to clarify whether this frequency-specific neural tracking is the result of intrinsic neural oscillations or stimulus-evoked responses (see a review, Obleser & Kayser, 2019). For instance, researchers could manipulate the speech rate (Oganian et al., 2023) and examine whether the frequency characteristics of neural tracking at the semantic level change in response to the varying word rates.

The present study has some limitations to be noted. First, there were several significant TRF responses to word entropy not included in the above analyses and discussions, which were primarily observed at the no-noise level. As the focus of the present study was speech-in-noise comprehension, these responses were not further discussed. Nevertheless, they also reflected speech information processing that would deserve investigations in future studies. Second, while the present study only adopted word entropy and word surprisal as two semantic-level features (Gillis et al., 2021; Goldstein et al., 2022; Heilbron et al., 2022; Weissbart et al., 2020; Willems et al., 2016), the rapid development in NLP methods especially the large language models (LLMs) present us with a broader range of options such as semantic embedding (Heilbron et al., 2022). Future studies could employ additional indexes to fully demonstrate the adaptation mechanism of speech-in-noise comprehension. Furthermore, beyond feature extraction, the LLMs also could serve as brain-aligned agents which could be compared with humans and help unveil shared (or unique) mechanisms in the human brain (Caucheteux et al., 2023; Goldstein et al., 2022; Mahowald et al., 2023; Schrimpf et al., 2020). In sum, future studies could employ the promising NLP-based approach to further extend our understanding of language processing. Third, despite the advantage of the high temporal resolution of EEG in exploring temporal dynamics, the relatively poor spatial resolution limits the ability to investigate brain regions involved in predictive mechanisms. A more fine-grained analysis of the spatiotemporal dynamics of semantic prediction would require techniques such as fMRI, ECoG, or multimodal approaches.

In summary, the current study investigated how noise affected acoustic and semantic processing during naturalistic speech comprehension. With increasing noise, acoustic processing became increasingly delayed whereas semantic processing became increasingly advanced. Our results suggest that, while the efficiency of brain processing of speech information is indeed impaired by noise, the brain could compensate for the associated effects through active prediction at the semantic level. Overall, the present findings are expected to contribute to the growing research on the neural mechanisms of naturalistic speech comprehension in noisy environments.

## Funding

This work was supported by the National Natural Science Foundation of China (NSFC) and the German Research Foundation (DFG) in project Crossmodal Learning (NSFC 62061136001/DFG TRR-169/C1, B1), and the National Natural Science Foundation of China (61977041).

## Author contributions

**Xinmiao Zhang**: Conceptualization, Methodology, Formal analyses, Investigation, Data curation, Writing – original draft, Writing – review & editing. **Jiawei Li**: Conceptualization, Methodology, Investigation, Writing – review & editing. **Zhuoran Li**: Conceptualization, Methodology, Investigation, Writing – review & editing. **Bo Hong**: Writing – review & editing, Funding acquisition. **Tongxiang Diao**: Writing – review & editing. **Xin Ma**: Writing – review & editing. **Guido Nolte**: Writing – review & editing, Funding acquisition. **Andreas K. Engel**: Writing – review & editing, Funding acquisition. **Dan Zhang**: Writing – review & editing, Funding acquisition, Supervision.

## Declaration of competing interest

All the co-authors declare that they have no conflict of interest.

## Data availability statement

Data in the current study are openly available in the Open Science Framework at https://osf.io/de9p6/, reference number DOI 10.17605/OSF.IO/DE9P6.

## Acknowledgment

The authors would like to thank Prof. Xiaoqin Wang for providing the shielded room for the experiment. The authors would like to thank Prof. Zhiyuan Liu and members of his lab for computing the NLP models. The authors would like to thank Lingyi Tao and Xuelin Wang for their help with data collection.

## 5 Supplementary materials

**Figure S1.**
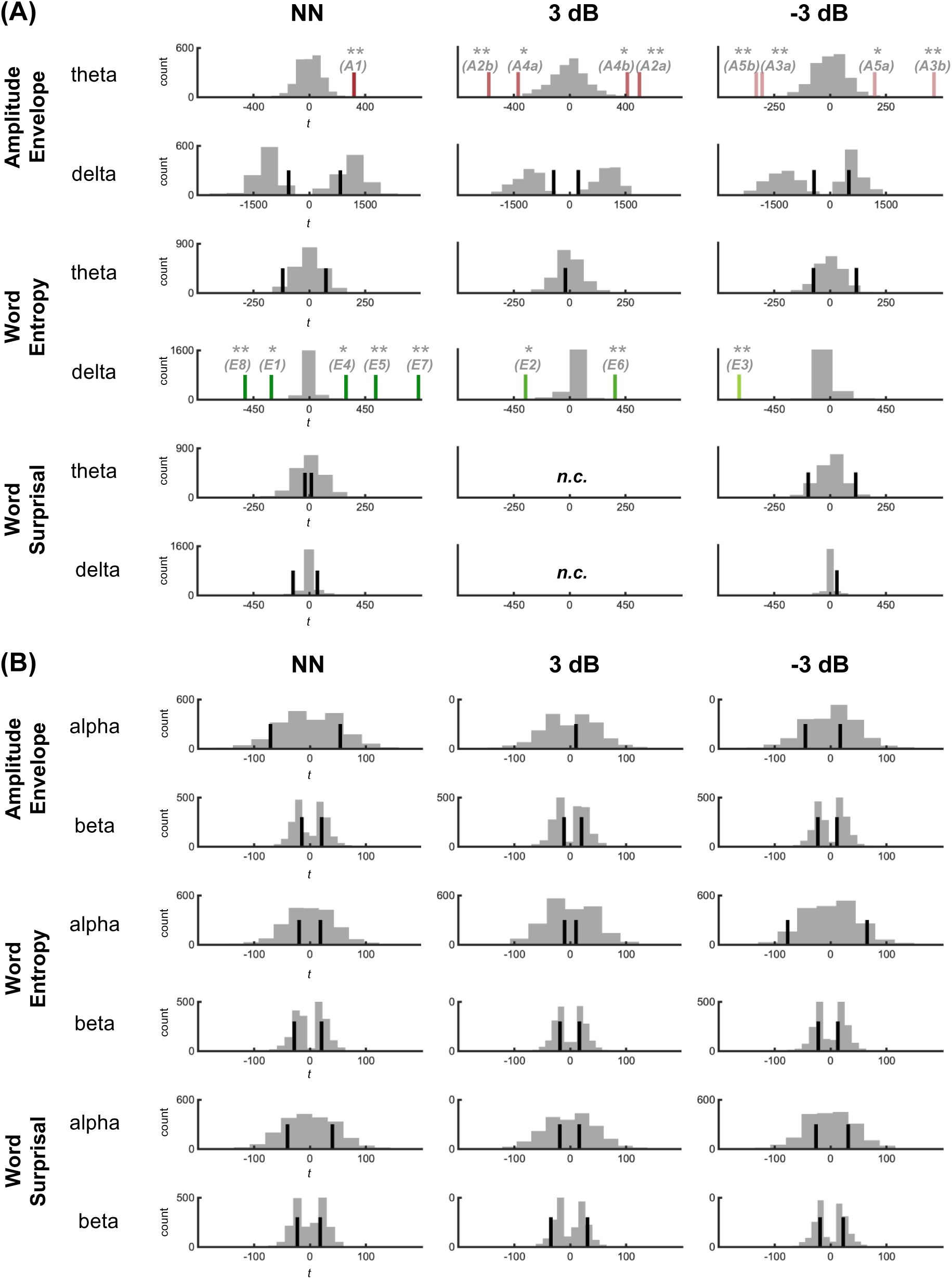
Clusters in TRF responses for different speech features at different SNR levels in the (A) delta/theta and (B) alpha/beta bands. The grey histograms show the distribution of the cluster-level test statistics from 1,000 permutations. The colored lines indicate significant clusters, and the black lines indicate nonsignificant clusters. n.c.: no cluster is formed. *: *p* < .05, **: *p* < .01, ***: *p* < .001.

**Figure S2.**
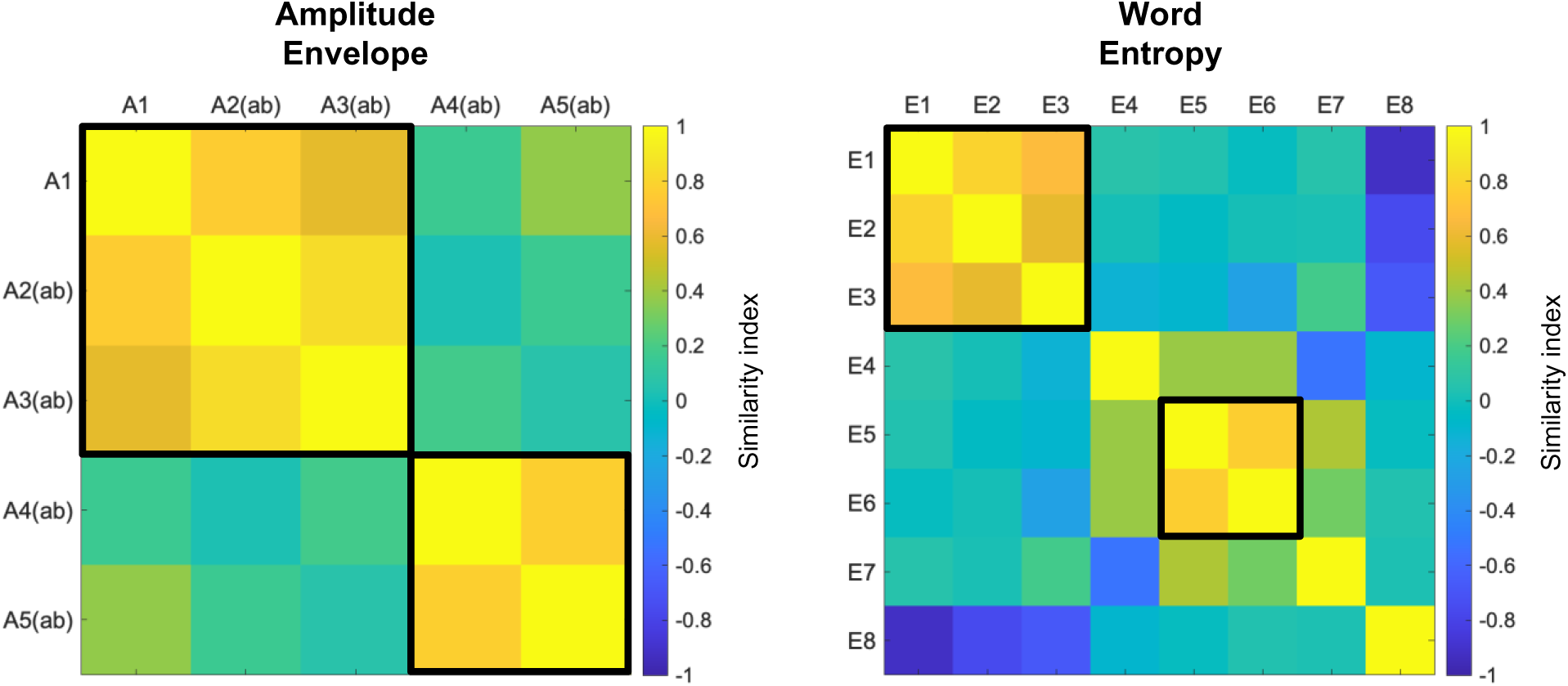
Similarity index of acoustic-level and semantic-level clusters. The similarity index was calculated as the following formula:

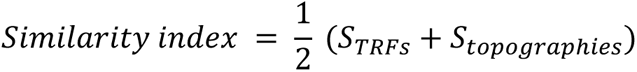

Where 𝑆_𝑇𝑅𝐹𝑠_ refers to the temporal similarity, which derived from the Pearson’s correlation between the averaged TRFs within the ROIs, and 𝑆_𝑡𝑜𝑝𝑜𝑔𝑟𝑎𝑝ℎ𝑖𝑒𝑠_ refers to the spatial similarity, which derived from the cosine similarity between topographies of the peak of clusters. Black rectangles indicate the visually identified clusters with similar spatiotemporal patterns.

**Figure S3.**
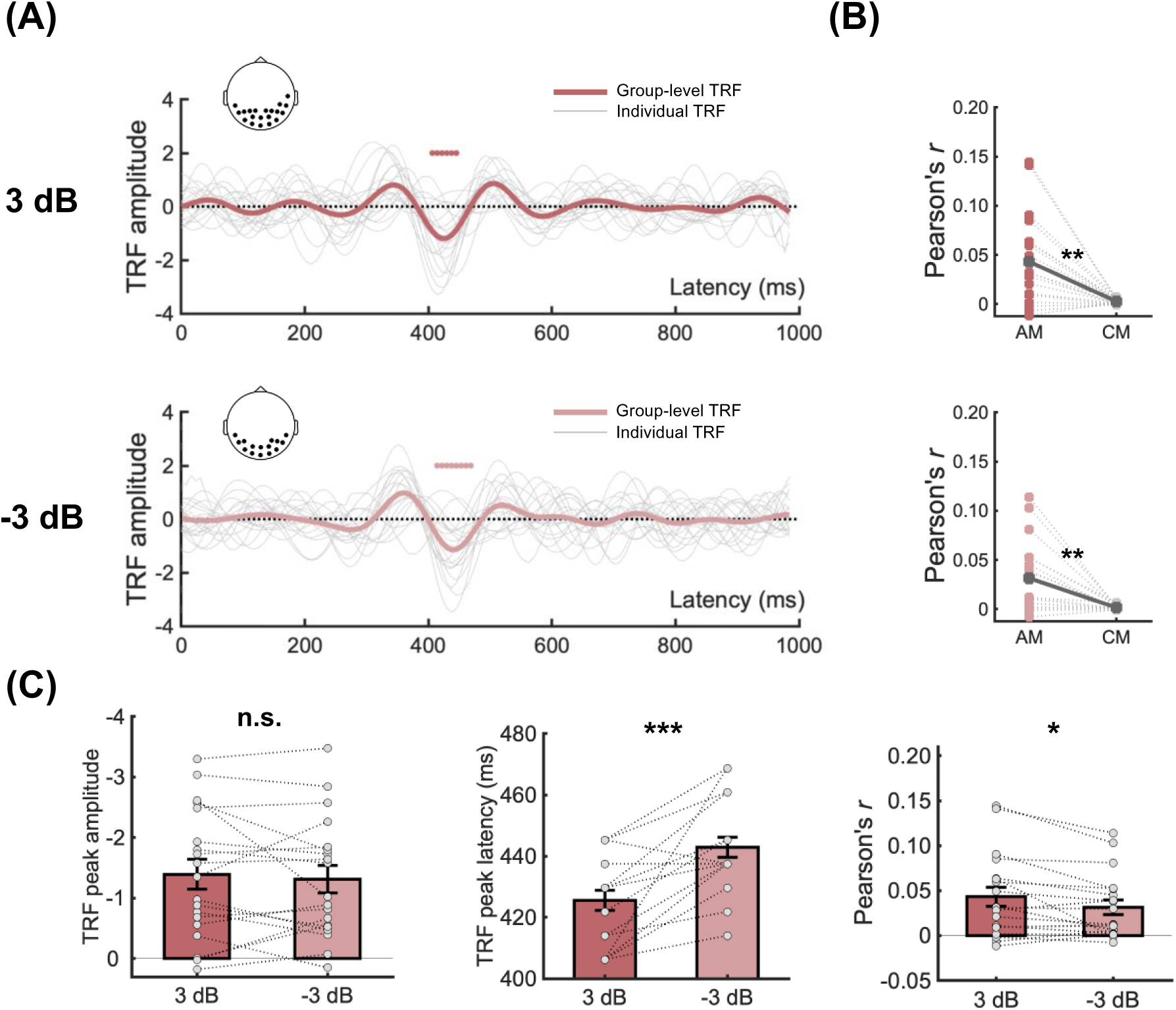
Cluster A4a and A5b of the acoustic-level TRF responses in the theta band. (A) The bold curves in different shades of red are the mean of TRFs averaged among ROIs across participants at the low- and high-noise levels. The grey curves are TRFs averaged among the ROIs of each participant. The colored horizontal line at the bottom of each plot indicates TOIs over which TRFs differed significantly from the control models. Dots in the corresponding topographies depicted the ROIs. (B) Reconstruction accuracy calculated from the ROIs/TOIs in (A). AM means actual models. CM means control models. (C) Noise effect on the peak amplitude, peak latency, and reconstruction accuracy. Peak latencies were significantly longer at the high-noise level than that at the low-noise level (paired-samples *t*-test, *t*(18) = 4.72, *p* < .001). Reconstruction accuracies were significantly lower at the high-noise level than that at the low-noise level (paired-samples *t*-test, *t*(18) = 2.54, *p* < .05). No significant differences were found in the peak amplitudes (paired-samples *t*-test, *t*(18) = 0.60, *p* = .558). Grey dots indicate data points from each participant. Error bar denotes the standard error. n.s.: not significant, *: *p* < .05, **: *p* < .01, ***: *p* < .001.

**Figure S4.**
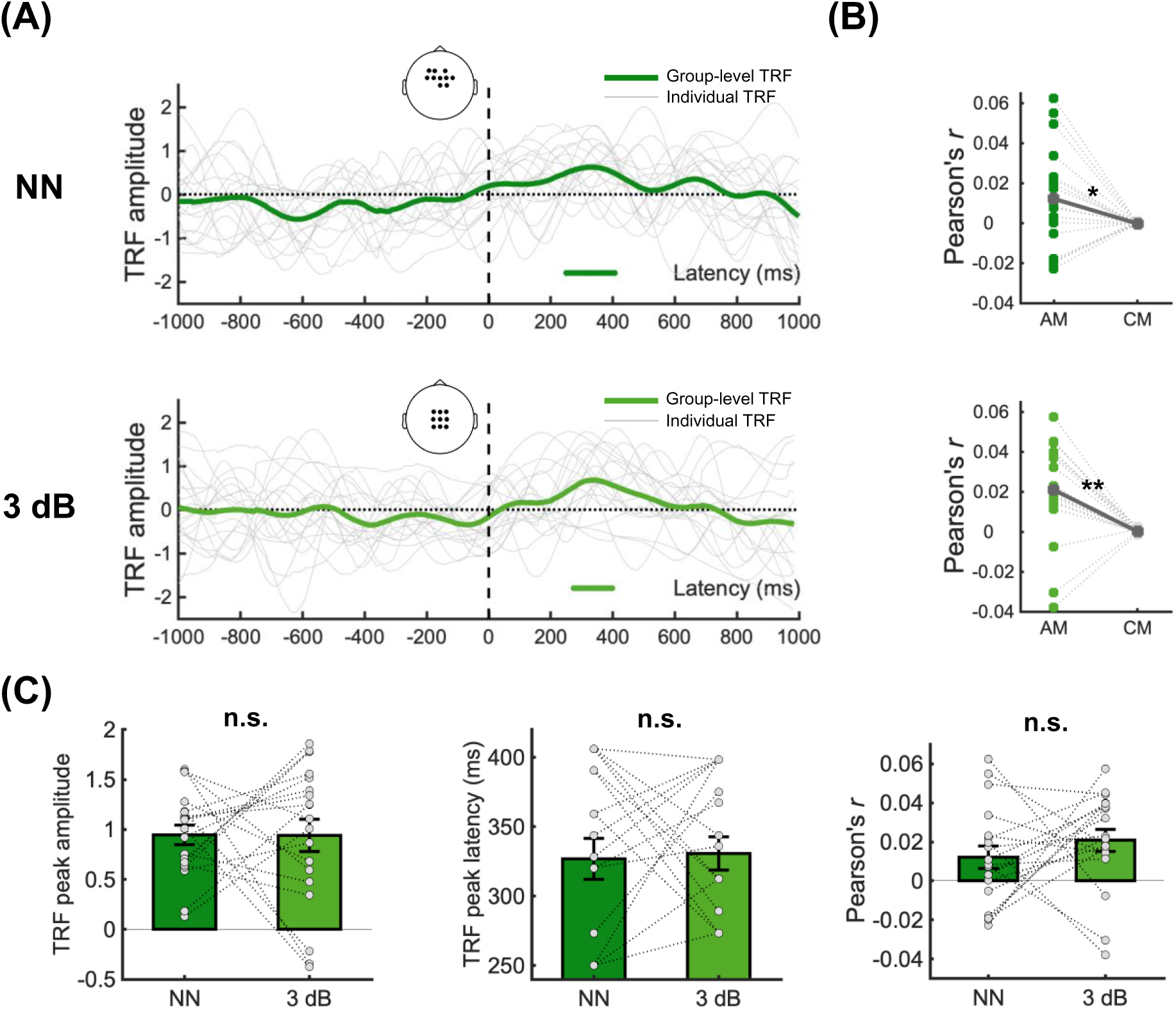
Cluster E5 and E6 of the semantic-level TRF responses to word entropy in the delta band. (A) The bold curves in different shades of green are the mean of TRFs averaged among ROIs across participants at the no- and low-noise levels. The grey curves are TRFs averaged among the ROIs of each participant. The colored horizontal line at the bottom of each plot indicates TOIs over which TRFs differed significantly from the control models. Dots in the corresponding topographies depicted the ROIs. (B) Reconstruction accuracy calculated from the ROIs/TOIs (A), AM means actual models. CM means control models. (C) Noise effect on the peak amplitude, peak latency, and reconstruction accuracy. No significant differences were found in the peak amplitudes (paired-samples *t*-test, *t*(18) = 0.02, *p* = .981), peak latencies (paired-samples *t*-test, *t*(18) = -0.21, *p* = .834), and reconstruction accuracies (paired-samples *t*-test, *t*(18) = 0.95, *p* = .357). Grey dots indicate data points from each participant. Error bar denotes the standard error. n.s.: not significant, *: *p* < .05, **: *p* < .01, ***: *p* < .001.

**Table S1.** Details of the Natural Language Processing model used to extract semantic features.

| Type | Parameter |
| --- | --- |
| Model type | Long-Short Term Memory (LSTM) |
| Embedding size | 200 |
| Hidden units per layer | 200 |
| Number of layers | 2 |
| Initial learning rate | 3 |
| Gradient clipping | 0.25 |
| Sequence length | 35 |
| Drop out | 0.2 |
| Epoch | 50 |
| Batch size | 3 |

**Table S2.** Descriptive statistics of semantic features. Std. means standard deviation.

|  | Number<br>of words | Word entropy |  |  | Word surprisal |  |  |
| --- | --- | --- | --- | --- | --- | --- | --- |
|  |  | Mean | Range | Std. | Mean | Range | Std. |
| **Story1** | 213 | 6.56 | 9.31 | 2.17 | 6.12 | 16.98 | 3.22 |
| **Story2** | 245 | 6.41 | 9.77 | 2.16 | 5.56 | 17.02 | 3.38 |
| **Story3** | 247 | 6.53 | 9.77 | 2.24 | 5.49 | 13.37 | 2.96 |
| **Story4** | 249 | 6.68 | 10.28 | 2.32 | 6.17 | 15.25 | 3.31 |
| **Story5** | 240 | 6.90 | 10.27 | 2.24 | 6.05 | 13.68 | 3.23 |
| **Story6** | 228 | 6.59 | 10.00 | 2.16 | 6.08 | 15.80 | 3.18 |
| **Story7** | 281 | 6.52 | 9.96 | 2.41 | 5.84 | 17.05 | 3.04 |
| **Story8** | 245 | 6.53 | 9.47 | 2.31 | 5.86 | 13.34 | 3.13 |
| **Story9** | 254 | 6.48 | 9.88 | 2.39 | 6.35 | 16.62 | 3.27 |
| **Story10** | 244 | 6.56 | 9.84 | 2.35 | 5.82 | 17.03 | 3.26 |
| **Story11** | 231 | 6.41 | 9.60 | 2.24 | 6.53 | 18.55 | 3.81 |
| **Story12** | 271 | 6.41 | 10.01 | 2.29 | 6.23 | 17.65 | 3.59 |
| **Story13** | 317 | 6.57 | 10.22 | 2.30 | 5.91 | 17.74 | 3.49 |
| **Story14** | 243 | 6.93 | 10.10 | 2.35 | 6.47 | 16.95 | 3.42 |
| **Story15** | 305 | 6.24 | 9.45 | 2.17 | 5.66 | 17.37 | 3.1 |
| **Story16** | 303 | 6.57 | 9.70 | 2.32 | 6.33 | 17.19 | 3.36 |
| **Story17** | 344 | 6.53 | 9.46 | 2.09 | 5.98 | 15.66 | 3.06 |
| **Story18** | 251 | 6.69 | 9.71 | 2.01 | 6.06 | 14.75 | 3.33 |
| **Story19** | 266 | 6.35 | 9.58 | 2.14 | 5.67 | 13.77 | 3.04 |
| **Story20** | 274 | 6.45 | 9.69 | 2.12 | 5.87 | 14.36 | 3.08 |
| **Story21** | 248 | 6.27 | 9.54 | 2.36 | 5.34 | 15.96 | 3.11 |
| **Story22** | 274 | 6.40 | 10.15 | 2.45 | 5.82 | 17.36 | 3.45 |
| **Story23** | 272 | 6.32 | 9.75 | 2.29 | 5.57 | 15.25 | 2.93 |
| **Story24** | 264 | 6.67 | 9.88 | 2.23 | 6.04 | 16.43 | 3.29 |
| **Story25** | 271 | 6.53 | 9.53 | 2.16 | 6.02 | 17.32 | 3.14 |
| **Story26** | 253 | 6.54 | 9.62 | 2.17 | 6.41 | 15.85 | 3.19 |
| **Story27** | 261 | 6.44 | 9.41 | 2.03 | 6.17 | 17.82 | 3.15 |
| **Story28** | 236 | 6.75 | 10.02 | 2.41 | 6.03 | 13.41 | 3.36 |
| **Story29** | 282 | 6.65 | 9.89 | 2.04 | 6.27 | 15.88 | 3.09 |
| **Story30** | 235 | 6.71 | 9.78 | 2.28 | 6.31 | 18.16 | 3.48 |

**Table S3.** Time lags of interests (TOIs) of each cluster in the acoustic-level TRF responses.

| Cluster | Time lags of interests (TOIs) |
| --- | --- |
| A1 | 383 ~ 483 ms |
| A2a | 406 ~ 453 ms |
| A2b | 484 ~ 539 ms |
| A3a | 344 ~ 383 ms |
| A3b | 414 ~ 469 ms |
| A4a | 406 ~ 445 ms |
| A4b | 484 ~ 531 ms |
| A5a | 336 ~ 383 ms |
| A5b | 414 ~ 469 ms |

**Table S4.** Time lags of interests (TOIs) of each cluster in the semantic-level TRF responses.

| Cluster | Time lags of interests (TOIs) |
| --- | --- |
| E1 | -297 ~ -195 ms |
| E2 | -398 ~ -250 ms |
| E3 | -625 ~ -359 ms |
| E4 | -8 ~ 141 ms |
| E5 | 250 ~ 406 ms |
| E6 | 273 ~ 398 ms |
| E7 | 602 ~ 961 ms |
| E8 | 563 ~ 727 ms |

**Table S5.** Spearman’s correlation between the reconstruction accuracies of each cluster in acoustic-level TRF responses and the behavioral performance. Significant correlation results are bolded (uncorrected).

|  | Comprehension<br>Performance | Clarity<br>Ratings | Intelligibility<br>Ratings |
| --- | --- | --- | --- |
| A1 | $r = .11, p = .647$ | $r = -.11, p = .662$ | $r = .10, p = .694$ |
| A2a | $r = .05, p = .831$ | **$r = -.46, p = .047$** | $r = -.04, p = .858$ |
| A2b | $r = .25, p = .309$ | **$r = -.50, p = .032$** | $r = .02, p = .926$ |
| A3a | $r = .09, p = .702$ | **$r = -.52, p = .025$** | $r = .05, p = .854$ |
| A3b | $r = -.002, p = .994$ | **$r = -.51, p = .028$** | $r = -.06, p = .798$ |
| A4a | $r = .08, p = .757$ | **$r = -.47, p = .043$** | $r = -.08, p = .734$ |
| A4b | $r = .18, p = .465$ | $r = -.42, p = .071$ | $r = .004, p = .986$ |
| A5a | $r = .03, p = .918$ | **$r = -.46, p = .048$** | $r = .002, p = .997$ |
| A5b | $r = .22, p = .361$ | **$r = -.52, p = .026$** | $r = .15, p = .541$ |

**Table S6.** Spearman’s correlation between the reconstruction accuracies of each cluster in semantic-level TRF responses and the behavioral performance. Significant correlation results are bolded (uncorrected).

|  | Comprehension<br>Performance | Clarity<br>Ratings | Intelligibility<br>Ratings |
| --- | --- | --- | --- |
| E1 | $r = -.09, p = .714$ | $r = -.21, p = .394$ | $r = .14, p = .560$ |
| E2 | $r = .11, p = .668$ | $r = .23, p = .339$ | $r = -.07, p = .762$ |
| E3 | $r = -.01, p = .997$ | $r = -.30, p = .206$ | $r = -.31, p = .203$ |
| E4 | $r = -.02, p = .928$ | $r = -.24, p = .332$ | $r = .09, p = .721$ |
| E5 | **$r = .51, p = .027$** | $r = .15, p = .527$ | $r = .37, p = .121$ |
| E6 | $r = .13, p = .594$ | $r = -.22, p = .365$ | $r = -.30, p = .215$ |
| E7 | $r = .20, p = .404$ | $r = .04, p = .881$ | $r = .15, p = .551$ |
| E8 | $r = .20, p = .411$ | $r = -.28, p = .237$ | $r = -.14, p = .575$ |

## Notes

### Competing Interest Statement

The authors have declared no competing interest.

### Summary of Updates

Updated results (minor revision for better visualization and improved clarity) and discussion.

